# Cross hybridization Inference for Phylogenetic Resolution (CIPHR)-FISH enables microbiome imaging with strain level taxonomic resolution

**DOI:** 10.64898/2026.02.26.708344

**Authors:** Emmanuel E. Adade, Ruogu Wang, Colin M. Henneberry, Alex A. Lemus, Rebecca J. Stevick, David Perez-Pascual, Bianca Audrain, Alexa J. Orsino, Dylan R. Farnsworth, Jean-Marc Ghigo, Alex M. Valm

## Abstract

The spatial organization of microbial communities is a critical determinant of host-microbe interactions, yet species-level mapping remains challenging due to high 16S rRNA sequence homology and spectral crosstalk in multiplexed fluorescence in situ hybridization (FISH). To address this challenge, we developed Cross-hybridization Inference for Phylogenetic Resolution (CIPHR)-FISH, a pipeline that integrates strategic probe design with supervised machine learning. CIPHR-FISH transforms probe cross-hybridization and spectral overlap, traditionally viewed as experimental noise, into informative molecular signatures. Using a gnotobiotic zebrafish model colonized with a defined mix of 10 zebrafish bacterial strains, we trained a support vector machine (SVM) on empirical hybridization patterns from pure bacterial cultures. CIPHR-FISH achieved 99.2 % macro-averaged accuracy, significantly outperforming standard linear unmixing (62.5 %), and successfully discriminated strains with 99.7% sequence homology. Applying this tool to gnotobiotic zebrafish larvae revealed distinct biogeographies: the intestinal bulb hosted highly structured, multi-layered polymicrobial aggregates, while the skin exhibited sparse, uniformly dispersed individual bacterial cells. Notably, we observed significant inter-individual variation in spatial community structure that was obscured by traditional bulk 16S rRNA sequencing. CIPHR-FISH provides a robust, scalable framework for high-resolution spatial biology by converting the limitations of molecular labeling into a rich data source for taxonomic classification. This approach enables the quantification of micro-scale ecological and stochastic forces that shape the microbiome across hosts.

## Introduction

The gut microbiomes of animals are extraordinarily complex communities composed of up to several thousand different species of bacteria, archaea and other microbes in e.g., humans (*1-3*). Just as the function of metazoan tissues and organs is mediated by the patterned assembly of phenotypically distinct cells during development, the functions of host-associated microbes in promoting health and disease arise from the emergent properties of their patterned structure(*1*, *3*) (*4*). The ever-decreasing cost of DNA sequencing has enabled a foundational understanding of the composition of the microbiome among healthy subjects, and changes that occur in community composition due to numerous factors, including diet and lifestyle(*5-10*). Attempts to move beyond compositional descriptions of human-associated microbial communities comprise an active area of bioinformatics research. High-throughput assays including metagenomics, metaproteomics and metabolomics are providing a path to investigate the interactions among gut microbes and between microbes and their host (*11-13*). Nonetheless, these techniques are typically applied to homogenized samples and neglect the spatial heterogeneity that underlies function within the complex metazoan gut ecosystem.

The gut community spatial structure, both biogeography and multicellular architecture, dictate host-microbe interactions and community functions (*14-17*). Along with changes in community composition, disruption of microbiome spatial structure is emerging as a hallmark feature of dysbiosis across host environments, especially with respect to the host epithelial mucus layer (*18-22*) (*23*).

Recent advances in multi-plex fluorescent imaging have demonstrated the ability to label and distinguish dozens of taxa of bacteria in practice and hundreds or up to 1000 different species, in principle, with state-of-the-art multispectral microscopy instrumentation, advanced computational image deconvolution, combinatorial labeling approaches, sequential labeling or other specialized fluorescence image acquisition methods (*24-27*). While in principle the number of distinct fluorescent readouts, i.e., spectrally variant fluorophores or, combinations thereof, approaches the number of species that comprise the gut microbiome, the ability to distinguish multiple species in a single community is greatly hindered by the extensive homology in 16S rRNA sequence among phylogenetically related species and strains (*28*). 18-20 mer single stranded DNA probes used in microbial fluorescence in situ hybridization are not discriminatory in the context of the single or few nucleotide differences that differentiate closely related bacteria in their 16S sequences (*29*).

To address the problem of species discrimination in 16S multiplex FISH, we introduce Cross-hybridization Inference for Phylogenetic Resolution (CIPHR)-FISH, a probe design, imaging and supervised learning-based image analysis tool. We hypothesized that while cross-hybridization among target species is unavoidable when using many species-specific probes in a single microbiome labeling scenario, small differences in hybridization efficiency would be observed when probes cross-hybridize to off-target species with nucleotide mismatches. Here the spectral crosstalk problem of multiplex fluorescence microscopy is convolved with probe cross-hybridization for species level 16S FISH. CIPHR-FISH learns the spectral cross-talk and probe cross hybridization patterns from images of labeled, lab-grown pure populations of the species of interest if they are available, as is the case with gnotobiotic animal models. To validate and benchmark CIPHR-FISH, we generated artificial mixtures of gut bacteria and employed a previously developed gnotobiotic zebrafish larva model, taking advantage of the availability of cultivatable bacterial isolates for model training, ease of generating axenic and gnotobiotic animals, the optical transparency of zebrafish larvae for imaging the gut and skin microbiota undisturbed within the gut, the richness of data generated from previous microbiome imaging in this model, and the known anatomical and functional parallels between zebrafish and higher animals (*30*-*32*) (*33*) (*34*, *35*).

## Results

We first manually designed FISH probes against each of the 10 strains in our gnotobiotic model such that each probe would have 100% on-target sequence homology and maximize the number of mismatches to all other, non-target strains in the consortium (*47*). With this logic, we identified 10 FISH probes that each had between 2-7 mismatches with all non-target strains in the community (**Figure 1A-B**), in which the 10 strains had between 70.7-99.7% homology in their 16S sequences (**Figure 1C**).

**Figure 1.**
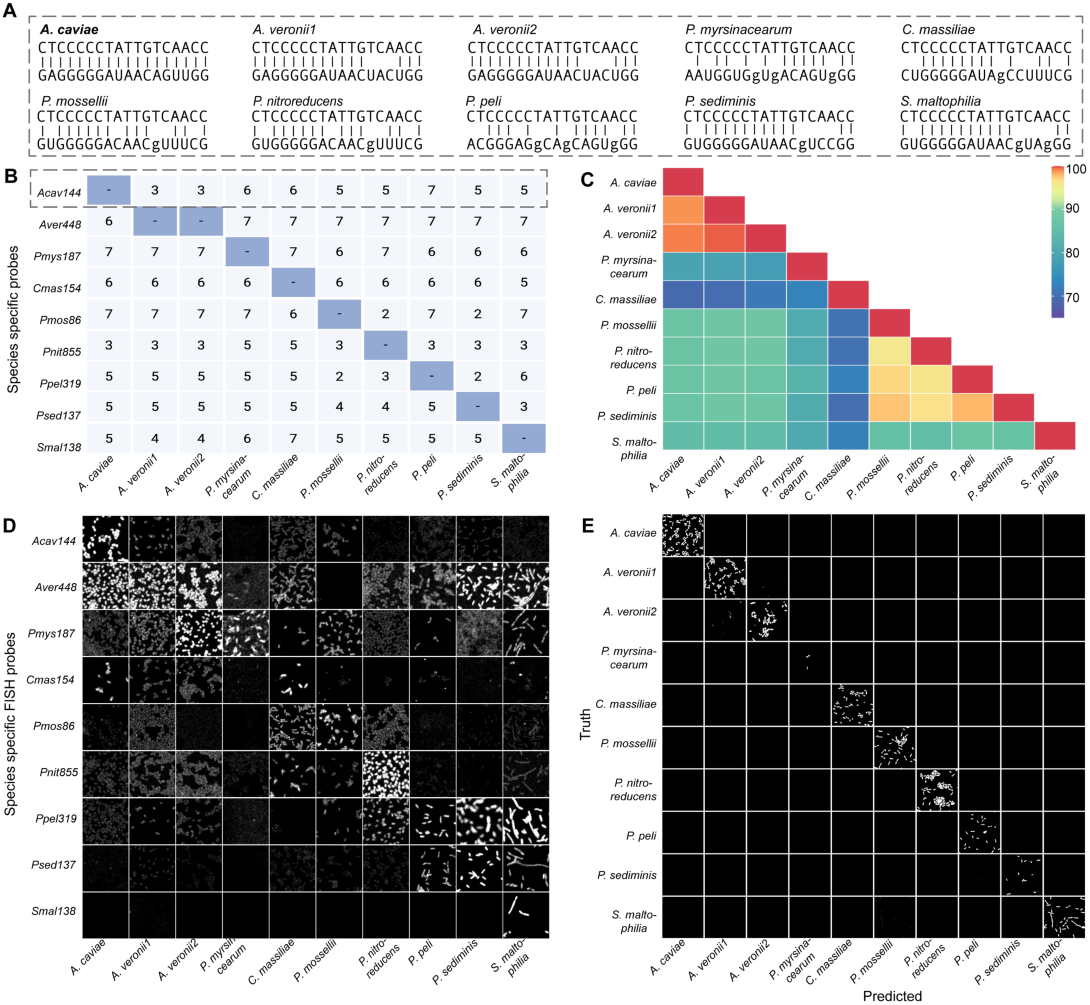
In silico design and evaluation of 16S rRNA FISH probes targeting closely related bacterial species from the zebrafish microbiome. (**A**) schematic of probe complementary and hybridization to on-target *A. caviae* and all other off-target species in our community, as a representative of all probe-target interactions. Asterisks indicate mismatches. (**B**) Minimum numbers of mismatches of each probe to all species in the community. - = 0 mismatches. (**C**) Sequence homology of the aligned 16S rRNA gene sequence among the community members. (**D**) Probe cross-hybridization matrix reveals distinct probe–microbial strain interaction signatures across microbial isolates with fluorescence signals reflecting probe hybridization to non-cognate and cognate targets. (**E**) Pixel-level classification of bacterial species and strains using a trained CIPHR model. Bar in (**D**) = 5 µm.

We had synthesized all 10 probes conjugated to the same fluorophore and performed quantitative imaging to assess probe specificity and cross-hybridization. We observed high specificity and variable cross-hybridization patterns for each probe and its target or non-target organisms. Visual inspection of the cross-hybridization matrix revealed a reproducible pattern in cross hybridization (**Figure 1D**). We next synthesized the 10 FISH probes, each conjugated to a unique, spectrally variant fluorescent reporter. We labeled pure populations of each of the 10 strains and acquired multi-spectral images of each population under identical acquisition settings. The resultant images served as labeled data for training a support vector machine (SVM) classification model, coded using the MATLAB Machine Learning Toolbox. We trained a classifier on a small 50 x 50 pixel subset image of each labeled species, that contained between 100-700 foreground pixels. We performed 5-fold cross validation using the Support Vector Machine Optimizable model with default setting of 30 iterations.

We benchmarked CIPHR-FISH against a state-of-the-art least squares-based linear unmixing algorithm (*48*). Using a pixel-wise argmax criterion (each pixel assigned to the channel with the highest estimated abundance), linear unmixing achieved a macro-averaged accuracy of 0.5747, an overall accuracy of 0.6249, and a macro F1 score of 0.5417 across four independent tests. In contrast, CIPHR achieved a macro-averaged accuracy of 0.9910, an overall accuracy of 0.9923, and a macro F1 score of 0.9856 across the same tests (**Table 1**). Here, macro-averaged accuracy is defined as the unweighted mean of per-taxon accuracies (each taxon contributes equally), whereas overall accuracy is computed by pooling all foreground pixels across cells (equivalently, a pixel-count–weighted average across species). The improvement of CIPHR over Linear Unmixing was consistent across species and statistically significant under a paired signed-rank test on per-taxon accuracies (p = 0.002) (**Table 2**). Visual inspection of images of pure populations after CIPHR-FISH classification demonstrate robust classification of species and strains as evidenced by their isolated appearance along the diagonal in the image matrix plot of species identity vs. CIPHR classification (**Figure 1E**). Lastly, we validated CIPHR-FISH on an artificial mixture of the 10 zebrafish isolates in which each bacterial strain varied in abundance by a one-fold increase from least to most abundant to simulate a complex community (**Figure 2B**). Output after CIPHR-FISH classification correlated with input into the mixed community with a coefficient of determination of 0.93 (**Figure 2C**).

**Figure 2.**
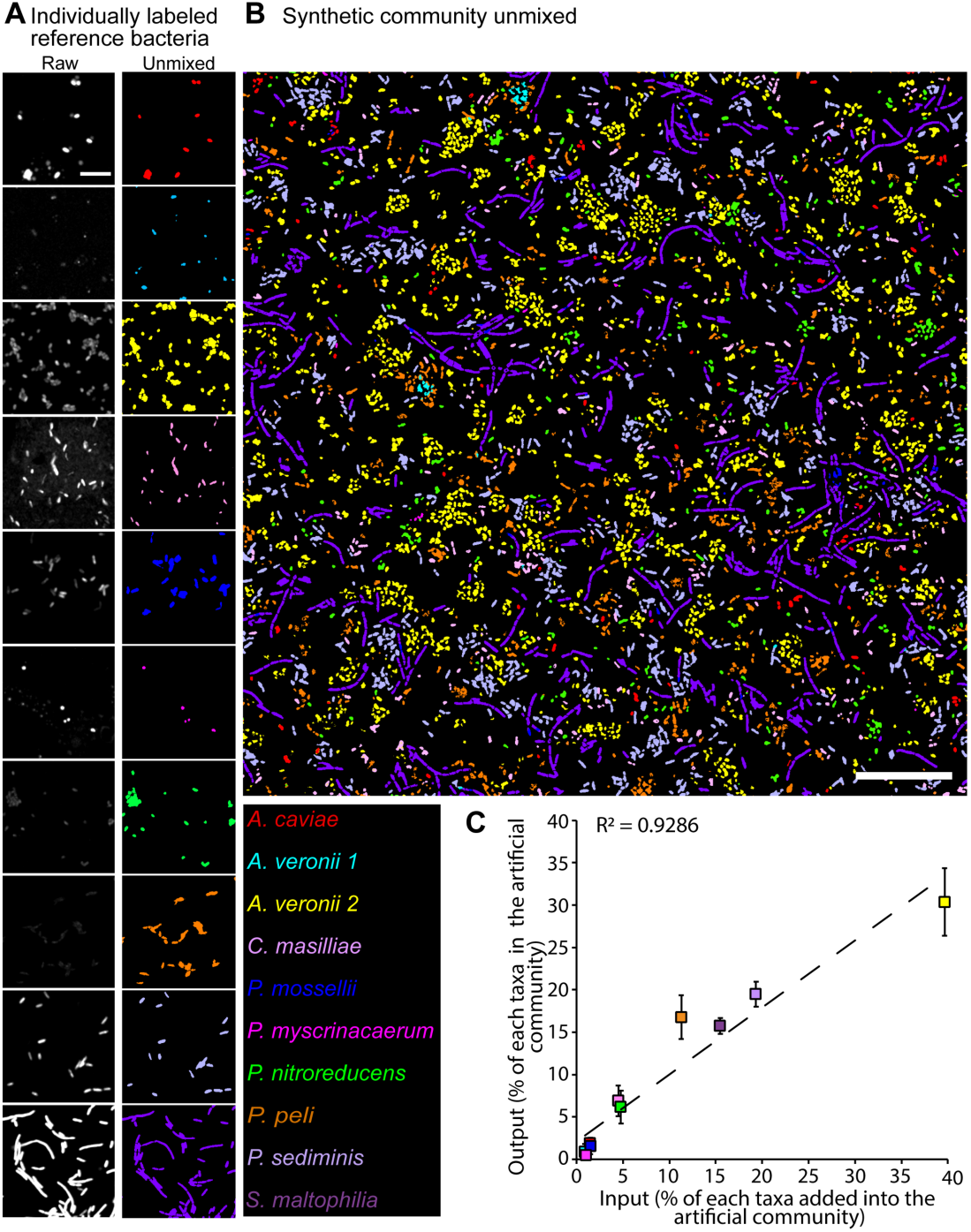
CIPHR-FISH imaging of pure bacterial populations and a 10-member defined community. (**A**) Individually labelled bacteria cells. Left images are raw images made from MIPs of all 82 concatenated spectral channels in gray scale and right images are pseudocolored images of the same field of view after CIPHR-FISH classification. Bar = 5 µm. (**B**) Representative CIPHR-FISH image of a defined bacterial community assembled from the same species in (**A)**, each in different proportion. Bar = 20 µm. (**C**) Plot of output from quantitative analysis of CIPHR-FISH images of the community in (**B**) verses input. Dashed line = expected output vs. input. Error bars = standard deviation from n = 4 images.

**Table 1.**
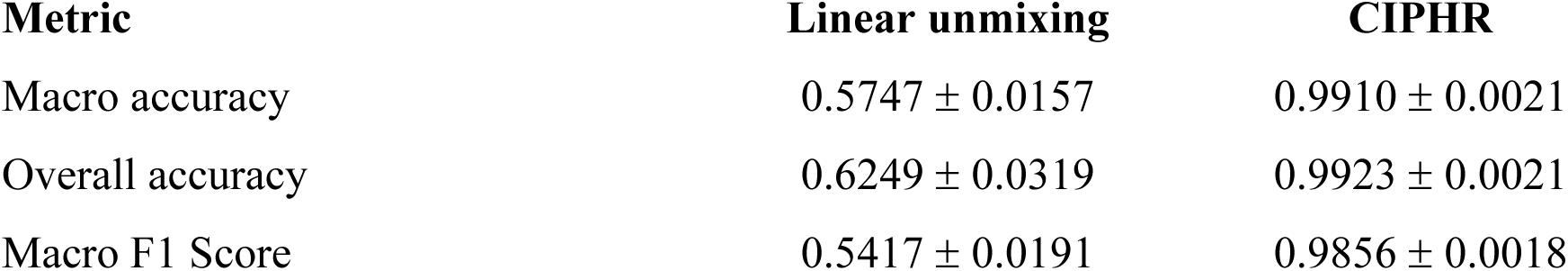
Method comparison summary of CIPHR-FISH classification to state-of-the-art linear unmixing.

**Table 2.** Quantitative comparison of CIPHR-FISH classification to state-of-the-art linear unmixing per taxon.

| **Bacterial species** | **Linear unmixing** | **CIPHR** |
| --- | --- | --- |
| *Aeromonas caviae* | $0.5792 \pm 0.0771$ | $0.9996 \pm 0.0001$ |
| *Aeromonas veronii 1* | $0.2355 \pm 0.0067$ | $0.9945 \pm 0.0035$ |
| *Aeromonas veronii 2* | $0.7664 \pm 0.0040$ | $0.9672 \pm 0.0280$ |
| *Phyllobacterium myrsinacearum* | $0.0867 \pm 0.0287$ | $1.0000 \pm 0.0000$ |
| *Chryseobacterium massiliae* | $0.9425 \pm 0.0261$ | $0.9996 \pm 0.0002$ |
| *Pseudomonas mosellii* | $0.5861 \pm 0.0484$ | $0.9957 \pm 0.0039$ |
| *Pseudomonas nitroreducens* | $0.5460 \pm 0.0736$ | $0.9939 \pm 0.0020$ |
| *Pseudomonas peli* | $0.2340 \pm 0.0324$ | $0.9873 \pm 0.0021$ |
| *Pseudomonas sediminis* | $0.7749 \pm 0.0944$ | $0.9969 \pm 0.0025$ |
| *Stenotrophomonas maltophilia* | $0.9952 \pm 0.0017$ | $0.9755 \pm 0.0140$ |

Analysis was performed on four independent CIPHR predicted test images and four independent linear unmixed images. Data represents mean precision with standard deviation.

We next applied CIPHR-FISH to real biological samples in the form of gnotobiotic zebrafish larvae colonized with a 10-member bacterial community (*36*). Species and strain-level imaging of the larval zebrafish gut across three animals revealed a patterned biogeography from bulb through the proximal and distal gut, with differences in species composition, spatial clustering, and total biomass (**Figure 3**, **Supplementary Figures 1-3**). We observed three types of architectures in the bulbs of each of the three animals, (1) a highly intermixed architecture composed of *Stenotrophomonas maltophilia, Aeromonas veronii 2* and *Pseudomonas mosselii* (**Figure 3C, C’**), (2) a highly structured community consisting of distinct layers with an inner core dominated by *Pseudomonas mosselii*, with *Aeromonas caviae and Pseudomonas peli* evenly and non-randomly distributed within, followed by a thick layer dominated by *Pseudomonas nitroreducens* with *Aeromonas caviae* evenly distributed throughout, then yet another layer consisting almost exclusively of *Pseudomonas peli*, then a monospecies layer of *Phyllobacterium myrsinacaerum*, and finally a most superficial and thin layer of *Aeromonas veronii (both strains 1and 2) and Aeromonas caviae*, in presumed close contact with the mucus layer of the epithelium (**Figure 3D, D’**). The third (3) architecture observed in the bulb consisted of two large monophyletic clusters in proximity, each consisting of *Pseudomonas peli* and *Aeromonas caviae* with other species arranged as small, intermixed clusters or single cells, especially in the distal portion of the bulb (**Figure 3E, E’**). In the proximal intestine, across all three animals we observed small intermixed, monospecies clusters. Interestingly, although the species compositions across the three animals were highly heterogeneous, the proximal gut communities were consistently different from those of the bulb in the same animal (**Figure 3F-H**). The distal intestine in all three animals had low biomass, with occasional small monospecies clusters present (**Figure 3I-K**). Total microbial biomass displayed a strong biogeographic trend across all animals being highest in the bulb, intermediate in the proximal gut and lowest in the distal gut. In contrast, the species composition and architecture were highly heterogeneous across the three animals (**Figure 3L-M**). Analysis of site preference revealed that some species displayed strong preference for one of the three gut sites, with high heterogeneity across the three animals (**Figure 3N**).

**Figure 3.**
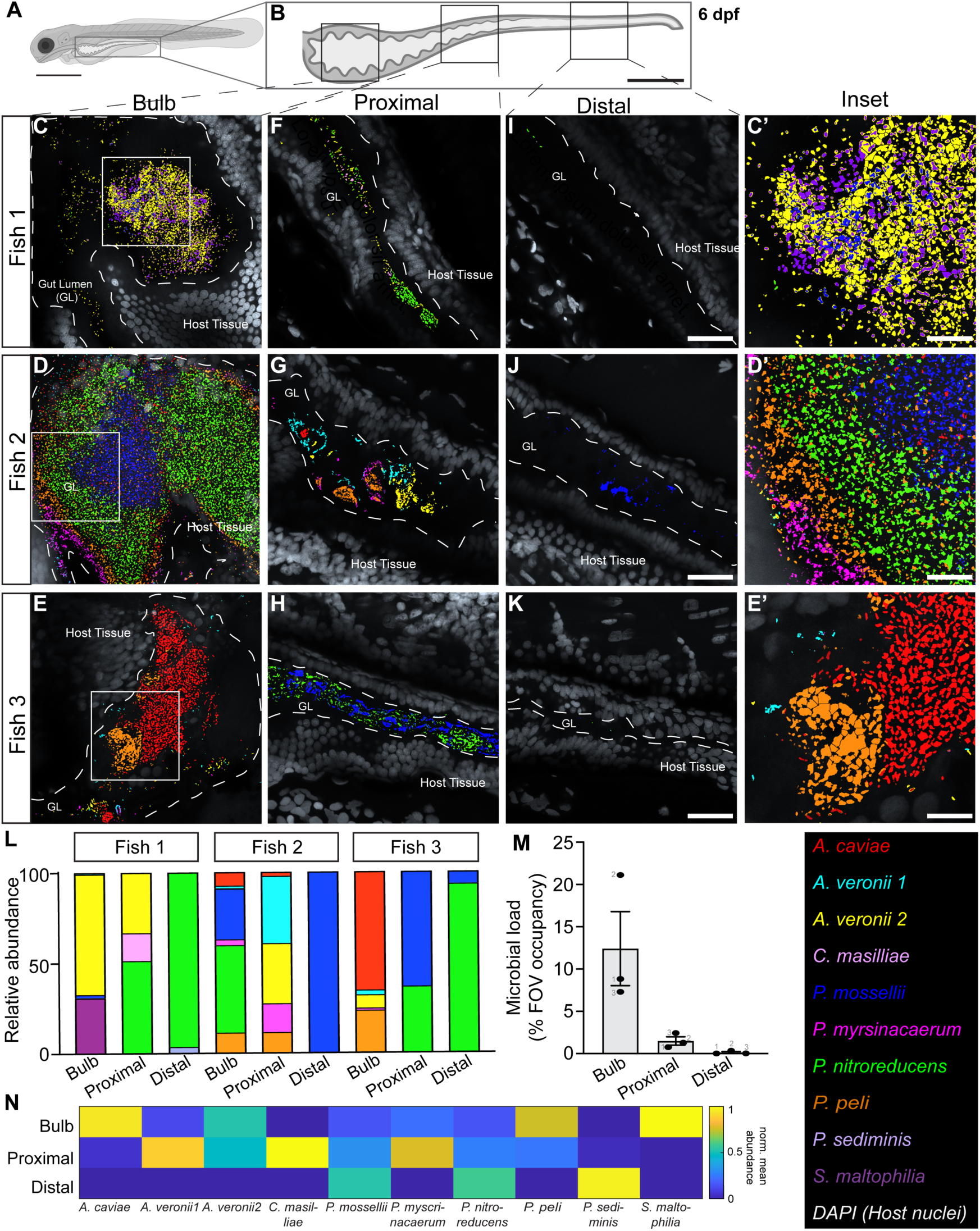
CIPHR-FISH imaging and analysis of gnotobiotic zebrafish gut. (**A**) Schematic of the larval zebrafish. Bar = 200 µm. (B) Schematic of zebrafish gut. Squares approximately delineate the three anatomical regions queried for spatial analyses of the gut microbiota, which we call bulb, proximal (intestine) and distal (intestine). Bar = 150 µm. Zebrafish schematics were generated in BioRender. (**C–K**) Representative images of the three gut regions after CIPHR-FISH image classification in each of 3 zebrafish larvae. Images are pseudocolored. Dashed lines indicate the epithelial border. Bars = 30 µm. (**C′, E′, H′**) Zoomed views of the boxed regions in (**C, E, H**). Bar = 5 µm. (**L**) Relative strain abundance plots of the three gut regions in each of the three larvae. (**M**) Mean total microbial biovolume measurements (n = 3 animals) in each of the three gut regions. Error bars represent the range of three biological replicates. (**N**) Heatmap of taxon-specific abundance within each gut region across three individual fish; data represent means from three biological replicates.

In contrast to the abundant clustering observed in the gut, the zebrafish skin microbiota exhibited a highly dispersed cellular architecture with some small aggregates (**Figure 4**). Multiple bacterial species were detected across animals in varying proportions; however, *Stenotrophomonas maltophilia* was consistently absent from the skin microbiota. Notably, one individual displayed a denser anterior microbial aggregate dominated by *Pseudomonas peli* and *Aeromonas caviae* near the gills (**Figure 4H**), whereas other fish displayed even, highly non-random dispersion of species as individual cells or small clusters across the skin. Biogeographic differences in microbial organization were observed along the anterior–posterior body axis. Skin-associated communities generally showed higher microbial densities in anterior regions with progressively reduced abundance toward the posterior. (**Figure 4K,L**). In contrast to the gut, most species present on skin did not display strong biogeographic preferences for any of the three ordinal skin regions that we designated here, suggesting consistent environmental habitats in the zebrafish skin apart from the region close to the gills (**Figure 4M**). Bulk 16S amplicon DNA sequencing of pooled animals in two separate experiments revealed strong similarity in microbiome composition among animals, in contrast to the highly heterogeneous imaging results obtained with CIPHR-FISH for three individual animals from the same experiment (**Supplementary Figure 4**).

**Figure 4.**
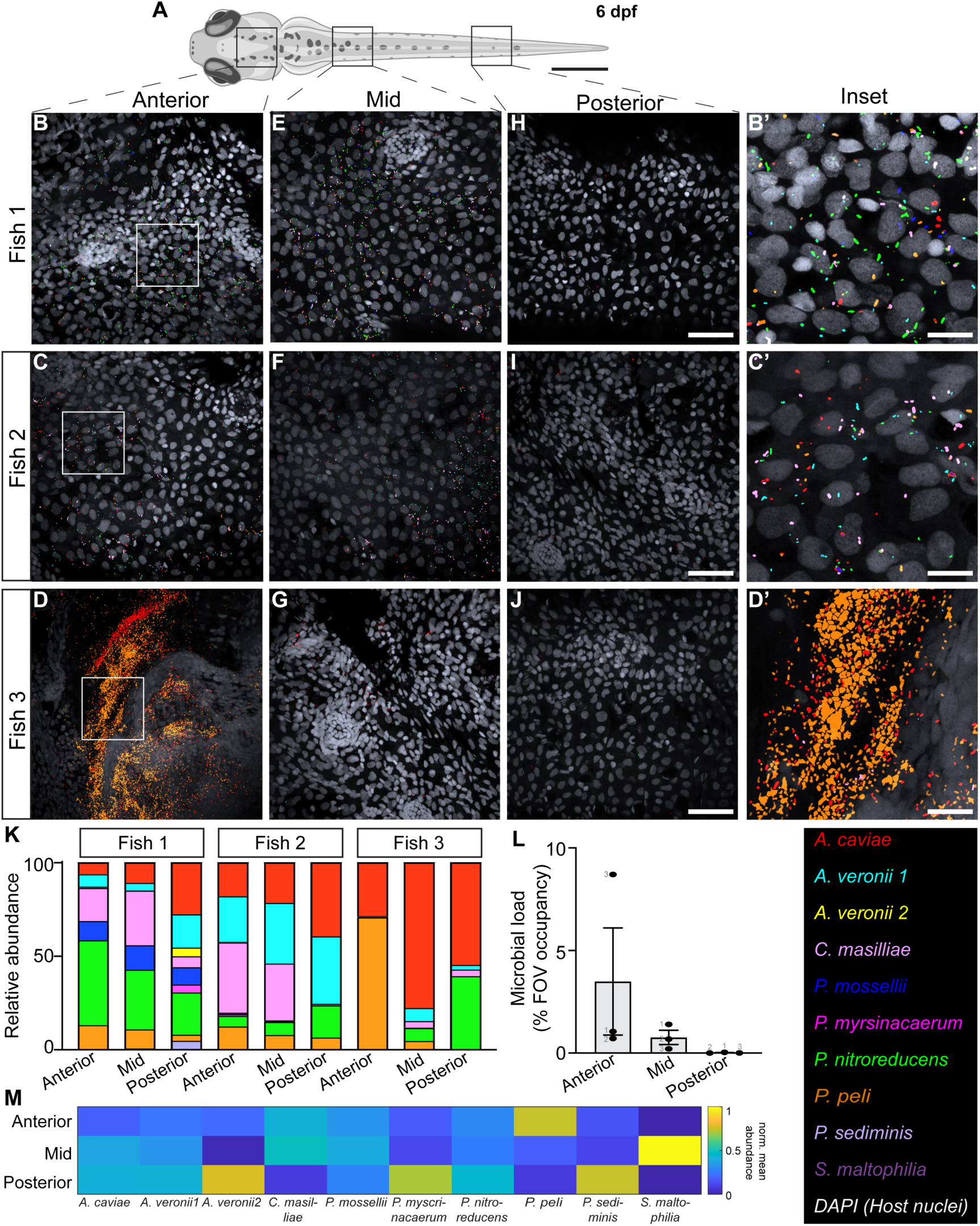
CIPHR-FISH imaging and analysis of gnotobiotic zebrafish skin. (**A**) Schematic of the larval zebrafish. Boxes approximate the three anatomical regions queried for spatial analyses of the skin microbiota. Bar = 200 µm. Zebrafish schematic was generated in BioRender. (**B–J**) Representative images of the three gut regions after CIPHR-FISH image classification in each of 3 zebrafish larvae. Images are pseudocolored. Bars = 30 µm. (**B′, E′, H′)** Zoomed views of boxed regions in (**B, E, H**). Bars = 7.5 µm. (**K**) Relative strain abundance plots of the three skin regions in each of the three larvae. (**L**) Microbial load measurements revealing region-specific distributions of species; error bars represent the range (minimum to maximum) of three biological replicates. (**M**) Mean total microbial biovolume measurements (n = 3 animals) in each of the three gut regions. Error bars represent the range of three biological replicates.

## Discussion

While in almost all cases it is possible, in principle, to discriminate bacterial species with specific 16S sequences due to the circular logic of using this gene as a phylogenetic marker; and, while on the order of 10^2^ spectrally variant bio-compatible fluorescent dyes are commercially available, the species-level mapping of complex microbial communities has been hampered by extensive probe cross-hybridization and spectral overlap of available dyes (*49*, *50*). In this study, we introduce CIPHR-FISH, a probe design and machine learning pipeline that transforms the problem of probe cross-hybridization and spectral crosstalk, traditionally treated as experimental artifacts to be avoided as much as possible, into informative molecular signatures that enable species-level resolution in microbial 16S FISH.

CIPHR-FISH also overcomes other limitations in microbial FISH. *In silico* predictions of probe hybridization efficiency often fail to accurately estimate probe brightness in labeled samples, most likely because of the difficulty in modeling tertiary structure and in predicting accessibility of the probes to regions of the 16S occluded by proteins in non-model organisms (*51-53*). In multiplex labeling applications, hybridization efficiency can vary significantly across probes because reaction conditions may be optimal for only one probe in the complex mixture, which may contribute to unexpected probe cross-hybridization (*54*). Heterogeneity in cell wall composition and thickness can also contribute to probe accessibility since cells are usually fixed by protein cross-linking agents (*55*). CIPHR-FISH elegantly transforms these probe and cell chemistry unknowns into valuable information that can be learned empirically from labeled cells and used to discriminate them in real biological samples. In fact, we demonstrated the ability to distinguish two strains of *Aeromonas veronii*, with 99.7% sequence homology. We cannot rule out that features other than sequence-based probe properties contributed to the unique fluorescent patterns that the CIPHR-FISH model used to distinguish these two strains.

Here we applied CIPHR-FISH to map the spatial biology of the gut and skin microbiomes of gnotobiotic zebrafish larvae. We observed, for the first time to our knowledge, the biogeography of a 10-member community of closely related species and strains in a gnotobiotic model. We observed dense polymicrobial aggregates in the gut, with multiple species-specific layers, especially in the bulb. This unexpected architecture suggests a strong ecological influence on community structure(*24*). Dense single species populations in lower complexity zebrafish models have been previously reported and unlike the highly acidic mammalian stomach, the zebrafish bulb is slightly alkaline at pH ∼7.5 (*56-59*). We further observed smaller monospecies clusters in the proximal intestine, similar to structures previously reported in murine models (*15*).

The skin of zebrafish larvae had a different community composition and biogeography compared to the gut, with most cells present as single cells or very small clusters. We cannot rule out that the method of reconventionalization that we employed, namely bath exposure to cultured microbial isolates during feeding, contributed to skin colonization; however, the highly non-random, uniform dispersion patterns of the microbiota and the complete absence of *S. maltophilia* from the skin, suggest an ecological and/or host anatomical influence on skin microbial biogeography (*60*). Interestingly, we observed high variance in microbial colonization patterns and community composition in individual gnotobiotic animals, a result that was not recapitulated with pooled DNA sequencing of the microbiota of animals from the same experiment, demonstrating the power of fluorescence imaging for studying micro-scale spatial structure and community heterogeneity in individual hosts.

While extraordinarily robust in species and strain identification in the scenarios to which we applied it here, CIPHR-FISH image analysis is a form of supervised learning that requires training on labeled data, i.e., cultivable isolates. While useful for in vitro studies and gnotobiotic animal models, we recognize that the SVM model that we employed here may not be appropriate for imaging complex microbial communities in conventional animals in which it is not practical or even possible to cultivate all microbiota species in the laboratory. In this case, CIPHR-FISH may be adapted to use semi- or unsupervised deep learning, such as with neural nets to be able to identify never-before-seen species from the perspective of the training model.

In summary, CIPHR-FISH comprises a scalable framework for species-resolved multiplex imaging by converting probe cross-reactivity into an informative readout for taxonomic classification. Application of this framework to animal models revealed that host-associated microbiota are governed by both region-specific ecological structuring and strong inter-individual variability. By unifying technological innovation with biological discovery, this work provides a foundation for future efforts to quantitatively map, understand, and manipulate host-associated microbial communities.

## Materials and methods

### Bacteria culturing and fixation

Bacterial species in the gnotobiotic model were originally isolated from zebrafish larvae and are listed in Supplementary Table 1 (*36*). All strains were grown in lysogeny broth (LB) (Corning) and incubated at 28°C with shaking or on LB agar plates. For model training and benchmarking, bacterial cells were harvested at mid-log phase and fixed in 4 % paraformaldehyde (PFA) at RT for 1.5 hours.

### Generation of the controlled bacterial community in gnotobiotic zebrafish

To generate the controlled bacterial community prior to imaging, the relative proportions of individual cells of the 10 species described in Supplementary Table 1 were estimated using FISH-based image quantification across four fields of view (FOVs) for pure isolates acquired with the Carl Zeiss LSM 980 with the 20x/ 1.4 NA objective. We then constructed the defined community with incremental, one-fold increase in cell abundance from one species to the next.

### Probe design and validation

Encoding probes were designed using the software tool, ARB (v6.0.4). Probe design parameters were set to 18–21 nucleotides in length with a GC content of 45–55%, and a minimum mismatch threshold of three mismatches to non-target sequences. Probe thermodynamic properties were evaluated using mathFISH, with acceptable probe–target hybridization free energies (delta G) ranging from −10 to −16 kcal/mol (*37*). For specificity testing and cross-hybridization analysis, probes were conjugated to FITC. For CIPHR-FISH labeling, encoding probes were appended with readout sequences at their 3’ ends (*27*, *38*). 10 taxon-specific readout probes were synthesized, each with a different fluorophore attached to its 5’ end (**Supplementary Table 2**). Probes were purchased from Integrated DNA Technologies). For analysis of 16S similarity, sequences were aligned using nucleotide BLAST (National Center for Biotechnology Information (NCBI)) or Clustal Omega (European Molecular Biology Laboratory (EMBL)).

### CIPHR-FISH labeling of pure bacterial isolates and defined community

Fluorescence in situ hybridization was carried out in 1.5 mL Eppendorf tubes. 2 µL of encoding probe was added to 98 µL of hybridization buffer [0.09 M NaCl, 0.02 M Tris pH 7.5, 0.01 % SDS, 20 % formamide] for a final probe concentration of 2 µM. Cells were incubated at 46 °C overnight (18-24 hrs). The cells were then washed in 100 μl of wash 1 [0.09 M NaCl, 0.02 M Tris pH 7.5, 0.01 % SDS, 20 % formamide] for 15 mins at 48 °C. To each tube, 98 μl of hybridization buffer was added along with 2 µL of readout probe (final concentration 2 µM) and tubes were incubated for 1 hr at 46 °C. Cells were washed in 100 μl of wash 1c[0.09 M NaCl, 0.02 M Tris pH 7.5, 0.01 % SDS, 20 % formamide] for 15 mins at 48 °C followed by 100 μl of wash 2 [0.09 M NaCl, 0.02 M Tris pH 7.5, 0.01 % SDS] for 15 mins at 48 °C. Wash 2 step was repeated for a total of 3 times. Cells were then suspended in 100 μl of 0.02M Tris. Cells were spotted onto an ultrastick slide (Goldseal) and incubated in a humid chamber for 2-4 hrs in the dark. Slides were dehydrated in an ethanol series and air dried in the dark. Samples were mounted in Prolong Gold and allowed to cure for at least 24 hours before imaging.

### Zebrafish husbandry

All experiments were carried out in accordance with animal welfare laws, guidelines, and policies and were approved by The University at Albany Institutional Animal Care and Use Committee. ABC wildtype zebrafish were maintained in a laboratory breeding colony in accordance to established protocols (*39*). Adult zebrafish were bred naturally in system water and fertilized eggs were transferred to 100mm Petri dishes with embryo medium. Embryos were allowed to develop at 28.5°C in embryo media and staged by hours post fertilization according to morphological criteria (*40*). Embryo media was prepared by adding 5 mL of stock A [175.0 g NaCl,7.5 g KCl,24.0 g MgSO4, 4.125 g KH2PO4,and 1.375 g Na2HPO4 per 2 L RO water], 1 mL of stock B [5.5 g CaCl2 per 400 mL RO water], and 1 mL of stock C [6.84 g NaHCO3 per 200 mL RO water] per 1 L of final solution, pH 7.2).

### Germ-free and microbial association

Germ-free derivation procedure was adapted from (*41*). All procedures were performed at 28°C under a laminar microbiological hood with single-use glassware. Fertilized eggs were kept in 25 cm^3^ vented flasks (Corning 430639) with 13 mL of filter sterilized embryonic media until 4 days post fertilization (dpf) (10 – 30 eggs/flask), transferred to new flasks after hatching at 4 dpf (8 – 15 fish/flask). Animals were maintained at 28 °C under 14/10 hour light/dark cycle through 6 dpf. At the end of the experiment, zebrafish larvae were euthanized with ice-water for about 30 mins before DNA extraction or fixation. Fish were fed with sterile larvae feed starting at 4 dpf as previously described (*36*). Following collection from timed spawning, embryos were rinsed and maintained under sterile conditions in sterile antibiotic embryo medium (ABEM-100 mg/ml ampicillin (Fisher, 243282), 5 µg/ml kanamycin (Fisher, 245484) and 250 ng/ml amphotericin B (MP Biomedicals, 195043). Embryo sterilization was performed using sequential immersion in 0.1% polyvinylpyrrolidone-iodine (PVP-I) followed by dilute sterile bleach (∼0.003%), with multiple sterile embryo medium rinses between steps. After sterilization, embryos were transferred aseptically into sterile flask with EM and incubated at 28 °C under a 14 h light/10 h dark cycle. Daily visual check of flask content was performed. At 4 dpf prior to microbial association, sterility check was performed by culturing 50 ul of the flask containing sterile EM on Luria broth (LB) agar. For microbial association, overnight cultures were prepared from freezer stock of the 10 different bacteria strains. Microbial cells were washed with sterile EM. The bacteria concentration was normalized and equal amount of bacteria were added to each flask to reconventionalize the germ-free larvae (*36*). Microbial exposure was simultaneous with first feeding.

### Zebrafish DNA extraction and sequencing

Whole zebrafish larvae were homogenized using a motorized pestle. Three to four larvae were pooled per extraction in 200 µL sterile filtered water. DNA was extracted using the DNEasy Blood and Tissue kit (Qiagen) following the manufacturer’s protocol, except samples were incubated with 100 µL of 10 mg/ml Lysozyme at 37 °C for 45 min. After adding 300 µL lysis buffer, the mixture was transferred to a bead-beating tube and mechanically disrupted with bead beating in Digital Disruptor Genie (Scientific Industries, Model SI-DD38) for 5 min at maximum speed (3000 RPM) setting. DNA yield was quantified using a Qubit high-sensitivity dsDNA fluorescence assay. Near full length V1-V9 16S rDNA Single Molecule Real Time (SMRT) sequencing was performed on the Revio system (Pacific Biosciences). Samples were prepared using Zymo Research’s “Quick-16STM Full-Length Library Prep Kit” using the following primers: 27f AGRGTTYGATYMTGGCTCAG and 1492r RGYTACCTTGTTACGACTT. Samples were prepared via 1-step PCR and were pooled by equal volume. A MagBead cleanup was performed on the pooled library which was then prepared for sequencing using the PacBio HiFi prep kit 96 with the SMRTbell barcoded adapter plate 3.01. Final libraries underwent the PacBio binding and cleanup steps using the Revio SPRQ Polymerase kit. Prepared libraries were sequenced on the PacBio Revio with a 12hr run time. SMRTbell barcoded adapters were removed on instrument during CCS. Post-CCS HiFi reads underwent secondary demultiplexing via Lima (v2.12.0). Generated BAM files from the secondary demultiplexing were converted to fastq.gz files using samtools (v1.17) and pigz (v2.6).

Analyses followed the PacBio HiFi-16S-Workflow (*42*). Sequences were imported to Qiime2 for analysis (*43*). Primer sequences were removed using Qiime2’s cutadapt plugin (*44*). Sequences were then denoised using Qiime2’s dada2 plugin (*45*). Denoised sequences were placed into a feature table detailing which amplicon sequence variants (ASVs) were observed in which samples, and how many times each ASV was observed in each sample. Identified ASVs were taxonomically assessed using the a curated database of the full-length 16S sequences of the 10 member community and the VSEARCH utility within Qiime2’s feature-classifier plugin (*40*, *45*). ASVs were then collapsed to their lowest taxonomic units, and their counts were converted to reflect their relative frequency within a sample.

### Zebrafish fixation

Whole larval zebrafish were fixed with Carnoy’s fixative [60% ethyl alcohol, 30% chloroform, 10% glacial acetic acid] for 2 hours at -20 °C at 6 dpf. Samples were then post-fixed in 4 % paraformaldehyde at 4 °C for 12 hours. Samples were stored in 100 % methanol for at least 24 hours before FISH labeling.

### CIPHR-FISH labeling of gnotobiotic zebrafish

FISH was carried out in 1.5 mL Eppendorf tubes. 2 µL of encoding probe was added to 98 µL of hybridization buffer [0.09 M NaCl, 0.02 M Tris pH 7.5, 0.01 % SDS, 20 % formamide] for a final probe concentration of 2 µM. Cells were incubated at 46 °C overnight (18-24 hrs). The cells were then washed in 100 μl of wash 1 [0.09 M NaCl, 0.02 M Tris pH 7.5, 0.01 % SDS, 20 % formamide] for 15 mins at 48 °C. To each tube, 98 μl of hybridization buffer was added along with 2 µL of readout probe (final concentration 2 µM) and tubes were incubated for 1 hr at 46 °C. Cells were washed in 100 μl of wash 1c[0.09 M NaCl, 0.02 M Tris pH 7.5, 0.01 % SDS, 20 % formamide] for 15 mins at 48 °C followed by 100 μl of wash 2 [0.09 M NaCl, 0.02 M Tris pH 7.5, 0.01 % SDS] for 15 mins at 48 °C. Wash 2 step was repeated for a total of 3 times. Cells were then suspended in 100 μl of 0.02M Tris. Fish were washed in 500 µl of wash 1 [0.09 M NaCl, 0.02 M Tris pH 7.5, 0.01 % SDS, 20 % formamide] for 30 mins at 48 °C followed by 1000 µl of wash 2 [0.09 M NaCl, 0.02 M Tris pH 7.5, 0.01 % SDS] for 30 mins at 48 °C then resuspended in 500 µl of 0.02 M Tris. Samples were incubated in 200 µl of 0.55 µM DAPI for 10 mins at RT. Samples were washed twice with 0.2M Tris buffer. Samples were mounted on Ultrastick glass slides. Fish were oriented so that the lateral surface of the larvae laid on the glass slide. Specimens were mounted in Slowfade anti-fade reagent (Thermofisher).

### CIPHR-FISH imaging

A Zeiss LSM 980 confocal microscope (Carl Zeiss) was used to acquire a multi-channel spectral image. The images were acquired with the Plan-Apo 63X/Oil DIC 1.4 NA objective in lambda mode using the 32-anode spectral detector to sample multiple 8.9 nm wavelength bands. Sequential excitation with 488 nm, 514 nm, 561 nm, 594 nm and 633 nm lasers were used to capture images, generating a total of 82 excitation/emission combination channels. 2-D images were acquired for bacteria references and the defined bacterial community. For zebrafish, 3D image Z-stacks were acquired with between 10 - 135 z-planes depending on the thickness of the anatomical site (skin, bulb, or intestine). DAPI channels in zebrafish samples were acquired separately with the 405 nm laser and merged with the CIPHR-FISH images after classification.

### Image pre-processing before CIPHR Classification

Image pre-processing was done in Fiji is just ImageJ (Fiji) v1.54p (*46*). To suppress high-frequency noise while preserving structural detail, a median filter (radius = 2.0 pixels) was applied to each channel prior to further processing. A maximum-intensity projection (MIP) was then generated from the concatenated stack for segmentation. Local adaptive thresholding was performed on the MIP using Otsu, Bernsen, or Phansalkar algorithms, selected based on image contrast and background characteristics, to produce binary masks for cell segmentation.

### CIPHR classification of bacterial communities

CIPHR-FISH classification was performed in MATLAB using the Classification Learner app in the Machine Learning and Deep Learning Toolbox. To train a Support Vector Machine (SVM), generated a reference spectra dataset from the images of labeled pure isolates. Each pixel in the reference data set was assigned a species label {1,2, … 𝐒}. For every pixel, we stored its 𝐂 channel spectral vector as a feature and its species ID as the target label. The feature training matrix was set as 𝐗 ∈ ℝ^N_train_ × C^, where 𝐍_train_ is the total number of labeled pixels across the entire training images, and each row is the unique pixel spectrum (derived from spectral variance profiling). The target vector contains the true species label for each pixel, 𝐲 ∈ {1,2, …, 𝐒}^N_train_^. We used Bayesian optimization over 30 iterations which captures non-linear decision boundaries in high dimensional datasets as well as hyperparameter tuning via five iterations of cross validation. For each new unlabeled pixel in an unknown image, we perform similar 𝐂 channel spectral vector extraction 𝐱 ∈ ℝ^C^, and fed into the previously trained SVM model for pixel classification and species prediction 𝐬 ∈ {1, …, 𝐒}. We then applied our trained classifier to the mixture of artificial communities or zebrafish images. For all classification, the image setting and pre-processing settings are kept the same for the references (training data) and the biological image that was classified (test image).

### Quantitative evaluation of linear unmixing performance on reference images

For each foreground pixel in the unmixed reference images, we queried the fluorophore channel with the highest estimated abundance and counted it as correct if this channel matched the expected on-target fluorophore for that taxon. Each analysis was performed on four independent FOVs of testing outcome per taxon generated CIPHR and Linear unmixing.

### Quantification of defined bacterial communities

Four images (nFOV = 4) of the bacterial community were acquired and analyzed with CIPHR-FISH. After species classification, the number of cells recovered per taxon was tabulated. We performed linear regression to assess the goodness of fit between known input and CIPHR-FISH generated output.

### Quantification of bacterial relative abundance and microbial load (FOV occupancy)

All analyses were performed in MATLAB R2021b and graphs and visualization done in GraphPad Prism 10 v10.6.0 and R. After CIPHR classification, each TIFF stack contained ten registered planes corresponding to each of the ten bacterial strains. FISH signal was segmented in FIJI/ImageJ (v1.54p) using channel-specific thresholding and particle analysis to quantify taxon-specific signal area. f abundance was quantified using area-based metrics rather than cell counts to account for signal overlap and dense bacterial aggregates. For each taxon *i* within region *r*, taxon area was extracted from segmented masks and aggregated across fields of view and imaging depths within each fish and region. Relative abundance was calculated as the proportion of taxon-specific area relative to the total microbial area in the same region:

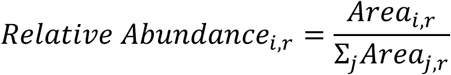

Where *Area_i,r_* represents the segmented FISH signal area for taxon *i* in region *r,* and the denominator represents the total microbial signal area across all cells in that region.

Microbial occupancy (% FOV coverage) was calculated as the ratio of the total microbial signal area to the total filed of view (FOV) area.

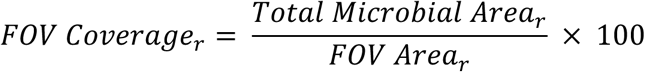

Where FOV area is determined by the image dimension.

### Spatial species distribution analysis

Heat maps were generated to visualize spatial taxon distribution across anatomical regions (gut: bulb, proximal, distal; skin: anterior, mid, posterior). Segmented FISH signal area was quantified and aggregated within fish, anatomical region and taxon level by summing taxon-specific area across all fields of view and imaging depths within each region, thereby avoiding pseudo-replication of technical measurements. Relative abundance was computed as the ratio of taxon area to total microbial area per fish and region, and mean values were then averaged across biological replicates (n = 3 fish) to construct taxon–region matrices. To emphasize spatial distribution patterns independent of overall abundance, matrices were row-normalized so that each taxon’s values summed to one across regions. Heat maps were generated in MATLAB.

## Supplementary Material

Supplementary material is included here.

## Supplementary information

**Supplementary Tables 1.**
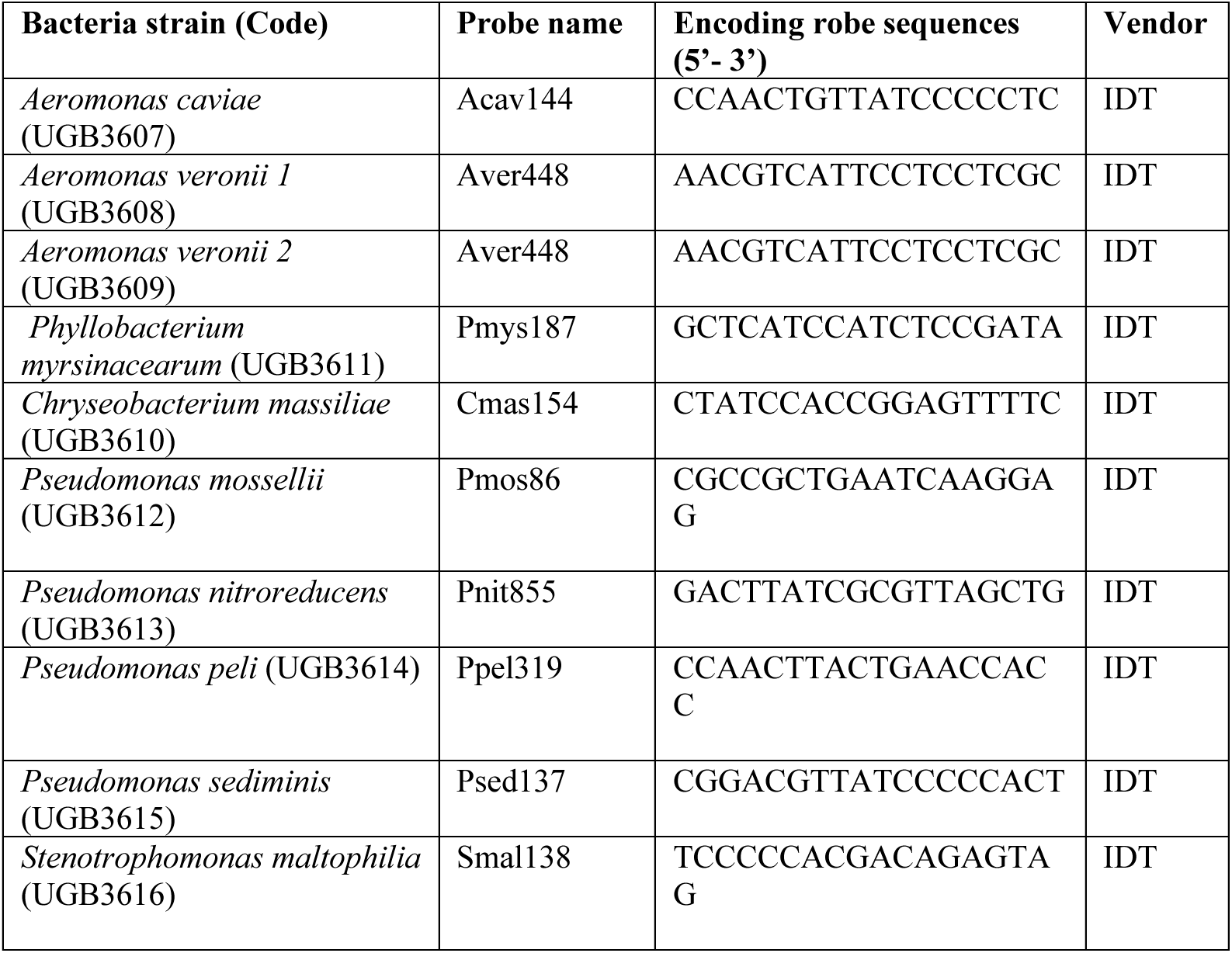
Salient features of bacterial strains and encoding FISH probes used in this study.

**Supplementary Table 2.**
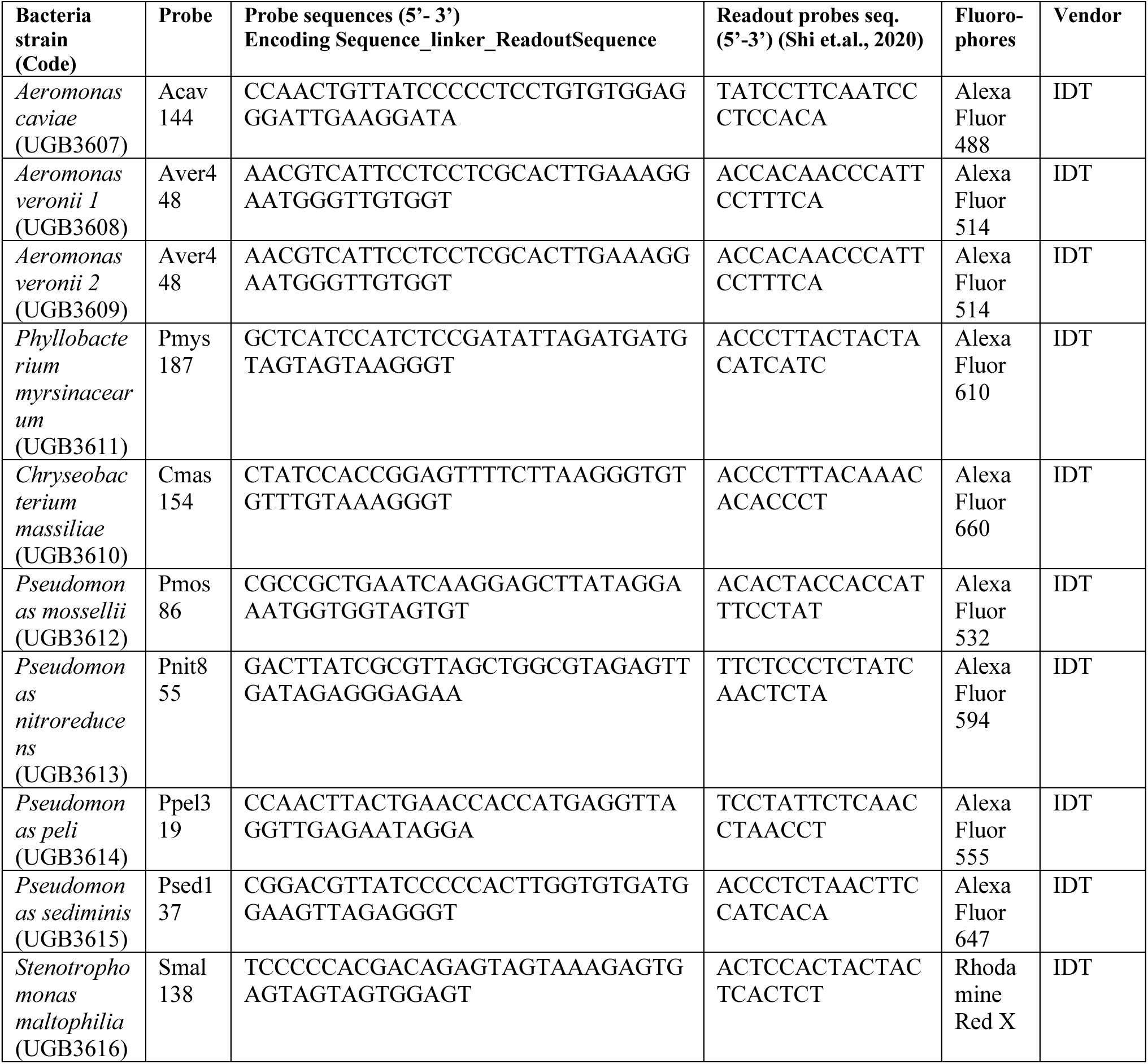
Encoding and Readout probe sequences and their conjugated fluorophores used in this study.

### Supplementary Figures

**Figure S1.**
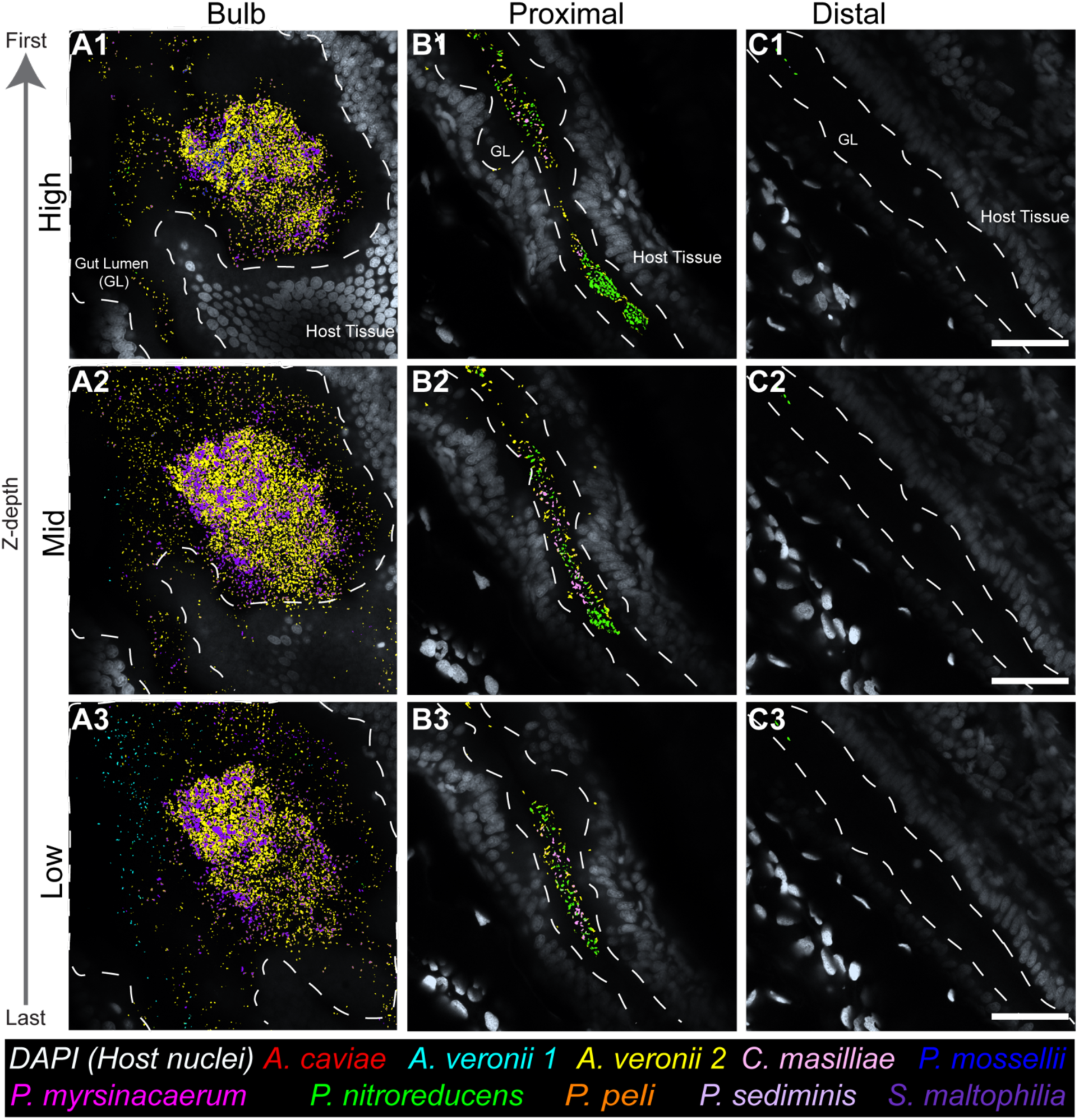
Species-level imaging of the larval zebrafish gut microbiome reveals spatially structured polymicrobial aggregates and internal variation across gut segments and through the axial depth of the gut in Fish 1. (**A1–A3**) Gut bulb biofilm architecture is dominated by *A. veronii*, with *S. maltophilia* and *P. mosselii* localized primarily within inner aggregate regions. (**B1–B3**) Proximal gut microbiome dominated by *P. nitroreducens* and *C. massilliae*. (**C1–C3)** Distal gut region predominantly colonized by *P. nitroreducens* with overall lower bacterial abundance. All panels show optical sections from confocal Z-stacks of species-resolved CIPHR-classified FISH signals from a defined 10-member microbial community. Bar = 30 µm.

**Figure S2.**
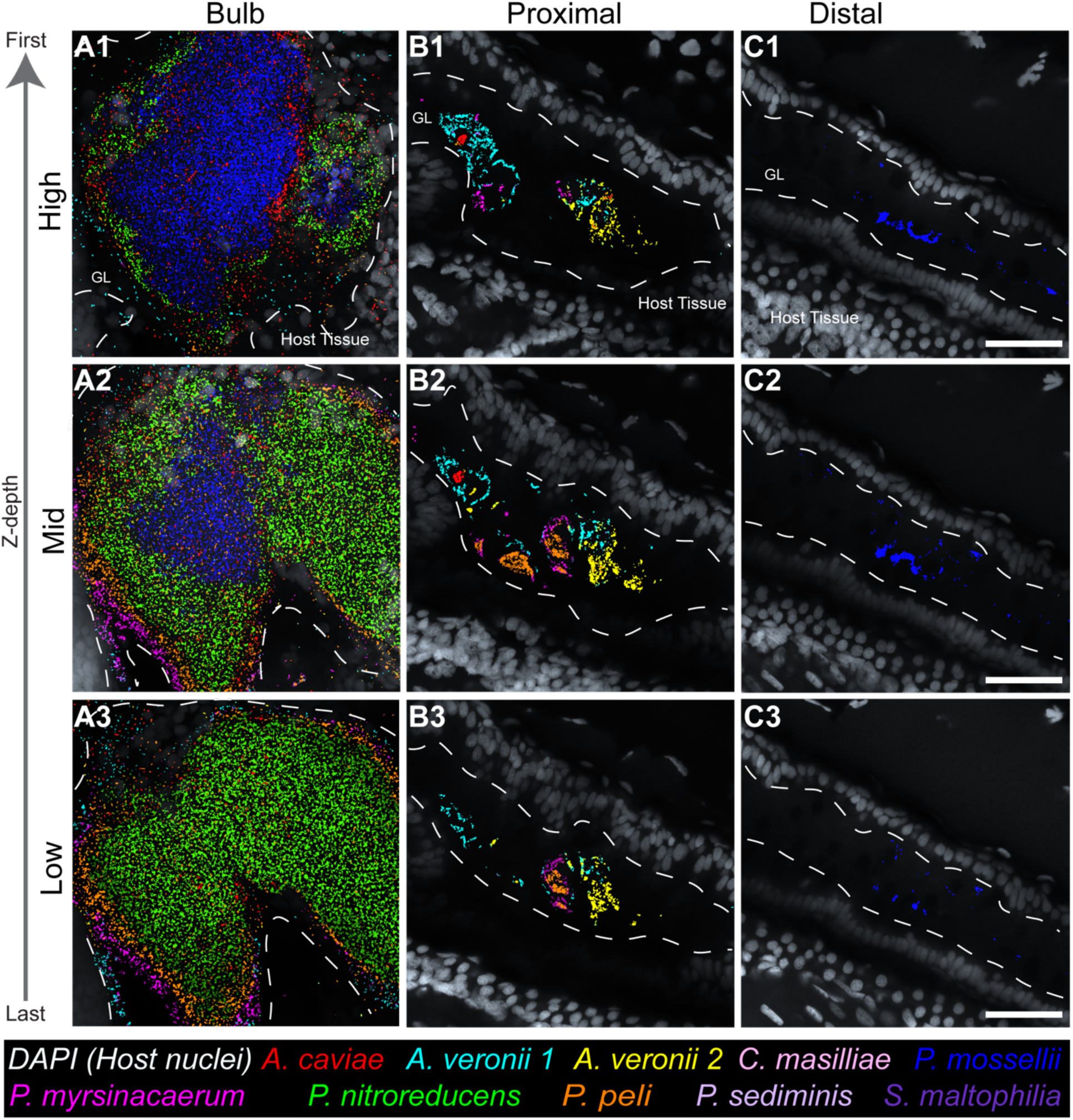
Species-level imaging of the larval zebrafish gut microbiome reveals spatially structured polymicrobial aggregates and internal variation across gut segments and along the Z-depth of Fish 3. (**A1–A3**) Gut bulb biofilm from a representative individual displays robust spatial organization. *Aeromonas veronii* (strains 1 and 2) and *A. caviae* localize near epithelial surfaces, while *P. myrsinacearum*, *P. peli*, and *P. mosselii* occupy outer aggregate regions. An inner core enriched in *Pseudomonas nitroreducens* and *P. mosselii*, with *A. caviae* distributed within the luminal biofilm space, is also observed (**B1–B3**) Proximal gut exhinited high microbial diversity with cells forming self-clustering micro aggregates (**C1–C3)** Distal gut region predominantly colonized by *P. mosselii,* with overall lower bacterial abundance. All panels show optical sections from confocal Z-stacks of species-resolved CIPHR-classified FISH signals from a defined 10-member microbial community. Bar = 30 µm.

**Figure S3.**
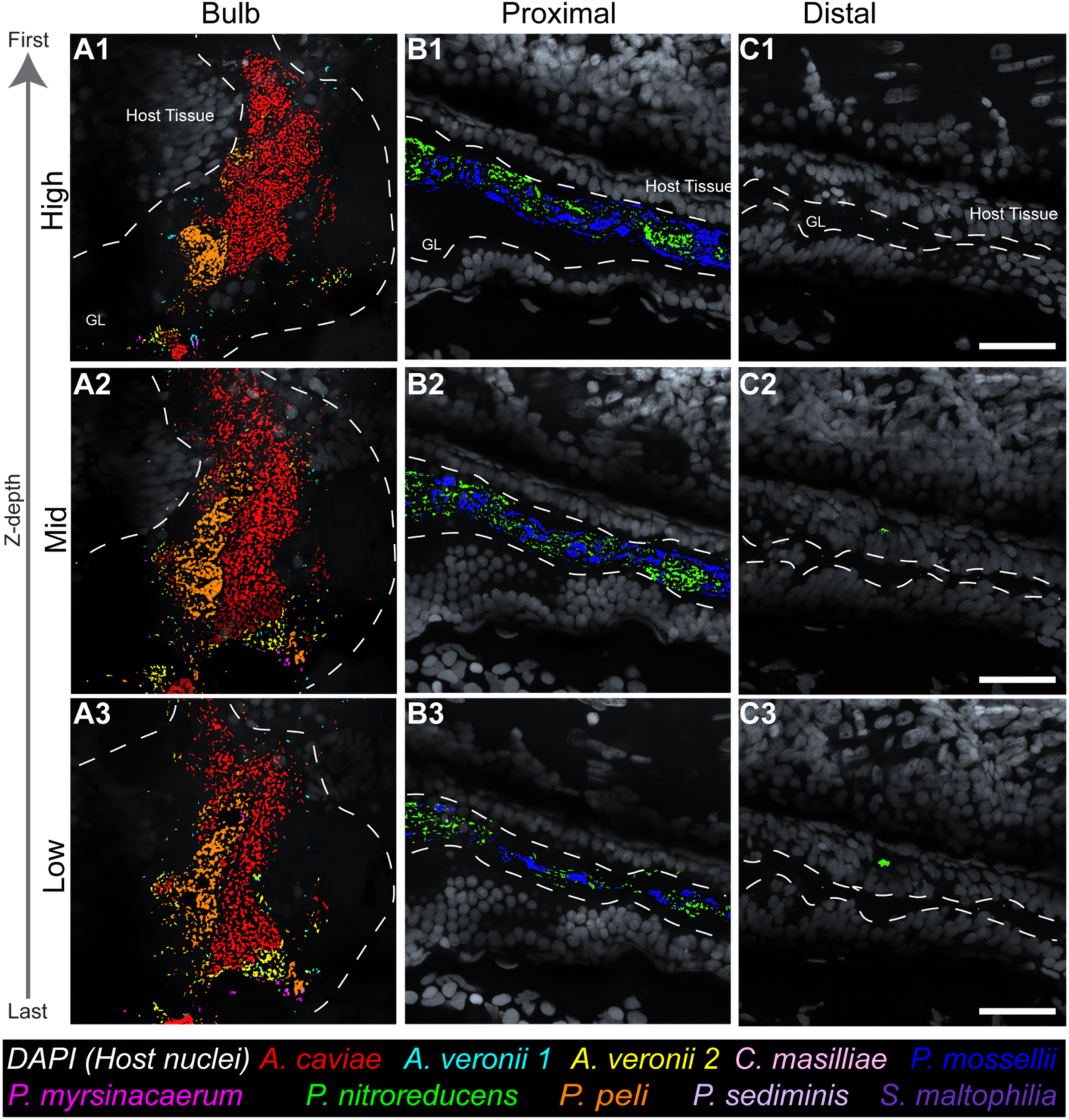
Species-level imaging of the larval zebrafish gut microbiome reveals spatially structured polymicrobial aggregates and internal variation across gut segments and along the Z-depth of Fish 2. (**A1–A3**) Gut bulb biofilm architecture dominated by *A. caviae*, with *P. peli* and *A.vaeronii* localized primarily within inner aggregate regions. (**B1–B3**) Proximal gut microbiome dominated by *P. nitroreducens* and *P. mossellli* in mono-species clusters (**C1–C3).** Distal gut region predominantly colonized by *P. nitroreducens,* with overall lower bacterial abundance. All panels show individual optical sections from confocal Z-stacks of species-resolved CIPHR-classified FISH signals from a defined 10-member microbial community. Bar = 30 µm.

**Figure S4.**
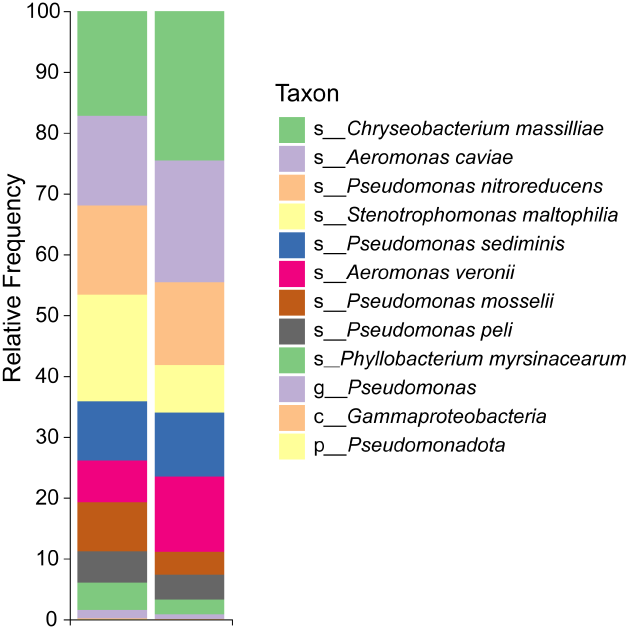
Full-length (V1–V9) 16S rRNA sequencing of reconventionalized zebrafish larvae colonized with a 10-member microbial consortium. DNA was extracted from whole zebrafish larvae in duplicate, with each replicate consisting of a pooled sample of four larvae to increase DNA yield.

